# CausalCell: applying causal discovery to single-cell analyses

**DOI:** 10.1101/2022.08.19.504494

**Authors:** Yujian Wen, Jielong Huang, Hai Zhang, Shuhui Guo, Yehezqel Elyahu, Alon Monsonego, Yanqing Ding, Hao Zhu

**Author notes:** Corresponding authors. (Y.D.), (H.Z.). These authors contributed equally to the work (Yujian Wen, Jielong Huang, Hai Zhang).

## Abstract

Correlation between objects does not answer many scientific questions because of the lack of causal but the excess of spurious information and is prone to happen by coincidence. Causal discovery infers causal relationships from data upon conditional independence test between objects without prior assumptions (e.g., variables have linear relationships and data follow the Gaussian distribution). Causal interactions within and between cells provide valuable information for investigating gene regulation, identifying diagnostic and therapeutic targets, and designing experimental and clinical studies. The rapid increase of single-cell data permits inferring causal interactions in many cell types. However, because no algorithms have been designed for handling abundant variables and few algorithms have been evaluated using real data, how to apply causal discovery to single-cell data remains a challenge. We report a pipeline and web server (http://www.gaemons.net/causalcell/causalDiscovery/) for accurately and conveniently performing causal discovery. The pipeline has been developed upon the benchmarking of 18 algorithms and the analyses of multiple datasets. Our applications indicate that only complicated algorithms can generate satisfactorily reliable results. Critical issues are discussed, and tips for best practices are provided.

## INTRODUCTION

The cell-specific regulation of gene expression and protein interaction generate various emergent signalling pathways which indicate that most interactions between genes and their products are causal. Causation determines widely observed and varied correlation. Some causal interactions are annotated in the “canonical” pathways (e.g., the KEGG pathways), but most remain unannotated, especially those in cells during development and in diseases and in small cell populations. On statistical data analysis, Judea Pearl wrote “*statistics alone cannot tell which is the cause and which is the effect*” (Pearl and Mackenzie, 2019); however, uncovering causation is more difficult than uncovering correlation. Causal discovery is a science which infers causal interactions from data observations upon testing conditional independence (CI) between variables (Glymour et al., 2019). Mathematically, CI is at the heart of causal discovery and CI≠unconditional independence ≠uncorrelation.

Researchers have used RNA-seq to detect gene expression in a lump of cells for years, but causal interactions in such mixed cells are blurred. Also, the sizes of such samples (lumped cells) are adequate for inferring only correlation but not causation between genes. Many methods (e.g., weighted gene co-expression network analysis, WGCNA) have been developed to construct networks of correlated genes in lumped cells upon RNA-seq data (Joehanes, 2018). Recently, scRNA-seq has been widely used to detect gene expression in single cells. In many situations (especially scRNA-seq using 10X Genomics), numbers of many cell types allow for inferring causal interactions between genes in each cell type.

Many different CI tests have been developed, from the quite fast Gauss CI test to the highly time-consuming kernel-based CI tests (Verbyla, 2018; Zhang et al., 2011). Gauss CI test is based upon partial correlations between variables; kernel-based CI tests estimate the dependence between variables upon their observations without assuming any relationship between variables or data distribution. CI tests critically characterize causal discovery algorithms and differentiate causal discovery from other network inference methods, including regulatory network inference (Nguyen et al., 2021; Pratapa et al., 2020), causal network inference (Lu et al., 2021), network inference (Deshpande et al., 2019), and gene network inference (Marbach et al., 2012).

The PC algorithm (named after its developers Peter Spirtes and Clark Glymour) is a state-of-the-art causal discovery algorithm and can work with different CI tests (Glymour et al., 2019). The time consumption of CI tests (especially kernel-based ones) makes it infeasible to apply the PC algorithm to all genes in a scRNA-seq dataset. On the other hand, what CI tests best suit scRNA-seq data and how to make proper trade-offs between time consumption and network size or accuracy remain unclear. Thus, benchmarking the PC algorithm and CI tests using single-cell data is essential before developing causal discovery pipelines and applying causal discovery to single-cell analysis.

This *Tools and Resources* article presents a solution to single-cell causal discovery by combining feature selection algorithms and causal discovery algorithms. Upon benchmarking 9 feature selection algorithms and 9 CI tests using simulated and real scRNA-seq data, we developed a pipeline and web server (called CausalCell) to perform causal discovery. Some measures are developed and imbeded into the pipelinle to ensure reliability of causal discovery. The analysis of multiple datasets were performed, with the results indicating that complicated (time-consuming) CI tests are crucial for generating reliable results. The inferred causal interactions provide informative clues for experimental and clinical studies.

## METHOD DESCRIPTION

### 1. Software implementation

The CausalCell pipeline consists mainly of feature selection and causal discovery. A parallel version of the PC algorithm (Le et al., 2019), together with the Docker techniques, is used to realize the parallel multi-task causal discovery, which is supported by a cluster of computers. The user interface is implemented using the Shiny language (Figure 1). Annotations of functions and parameters and a detailed description of an example are available online.

**Figure 1.**
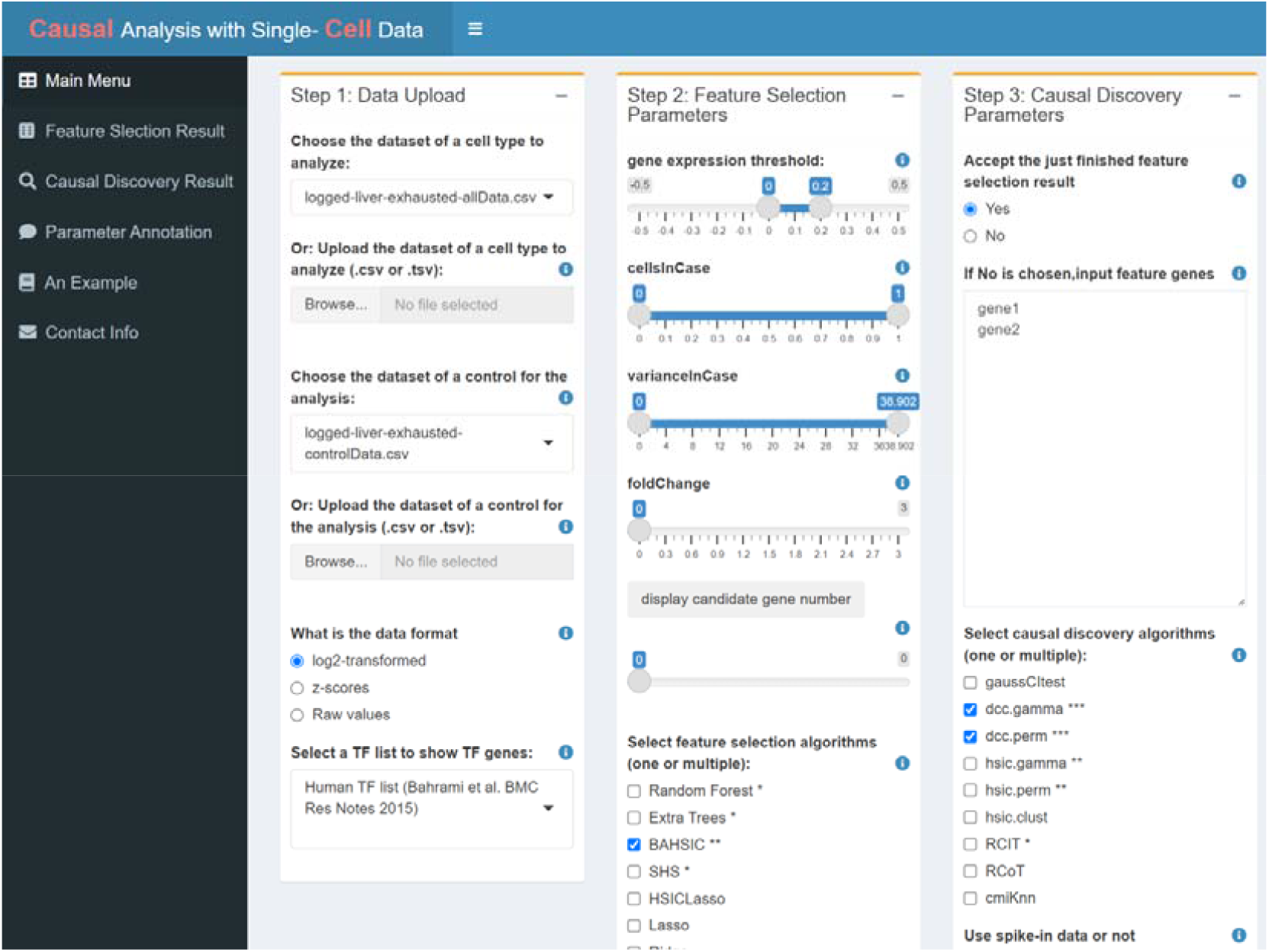
The user interface of CausalCell. Many functions are implemented to facilitate performing feature selection and causal discovery.

### 2. Data input and display

scRNA-seq data generated by multiple protocols (e.g., 10X Genomics, smart-seq2) and proteomics data (e.g., CyTOF) generated by mass cytometry can be analyzed (Supplementary Note 1). Data can be in the log2-transformed or z-score normalized format, and online transformation and normalization are available. A dataset can have or not have a control dataset. If a control dataset is uploaded, the fold change of gene expression is computed using the *FindMarkers* function in the *Seurat* package. Genes have multiple attributes (e.g., expression value, the percent of cells in which they are expressed, variance, and fold change); all of these attributes can be used to order genes to reveal gene expression features and to filter genes for performing feature selection (researchers often try to identify and analyse highly differentially expressed genes or genes having high variance).

### 3. Feature selection

Combining feature selection and causal discovery enables causal discovery to be applied to a arbitrary set of genes (feature genes). After genes are filtered upon conditions (i.e., expression threshold, the expressed cells, variance, and fold change) which generate candidate genes for feature selection, one or multiple genes of primary interest are used as the target genes (aka response variables) to select feature genes (aka features) from the candidate genes (aka candidates). Upon the evaluation of the accuracy, time consumption, and scalability of the 9 feature selection algorithms (Supplementary Note 2), BAHSIC is the most recommended feature selection algorithm. We also recommend the joint use of multiple algorithms (e.g., Random Forest + BAHSIC) to ensure reliability. Usually, feature genes should be 50-70 (depending on what causal discovery algorithms are chosen). Genes can be manually added into or removed from the feature gene list, to make feature genes better reflect a biological question. Also, all feature genes can be manually selected without performing feature selection, by which the user can examine any gene set.

### 4. Causal discovery

We implemented 9 causal discovery algorithms by combining the parallel version of the PC algorithm with 9 CI tests (Le et al., 2019). We evaluated the accuracy, time consumption, sample requirement, and stability of the 9 CI tests (Figure 2; Supplementary Note 3). The DCC algorithms are both most accurate and most time-consuming, suitable for small-scale network inferencec; RCIT is reasonably accurate and relatively fast, suitable for large-scale network inference. Multiple algorithms can be chosen in one run for a feature gene set, and a consensus network can be constructed upon the networks inferred by some or all selected algorithms. The consensus network is statistically more reliable. Edges in causal networks have arrows that indicate activation or inhibition and show thickness that indicate CI test’s statistical significance.

**Figure 2.**
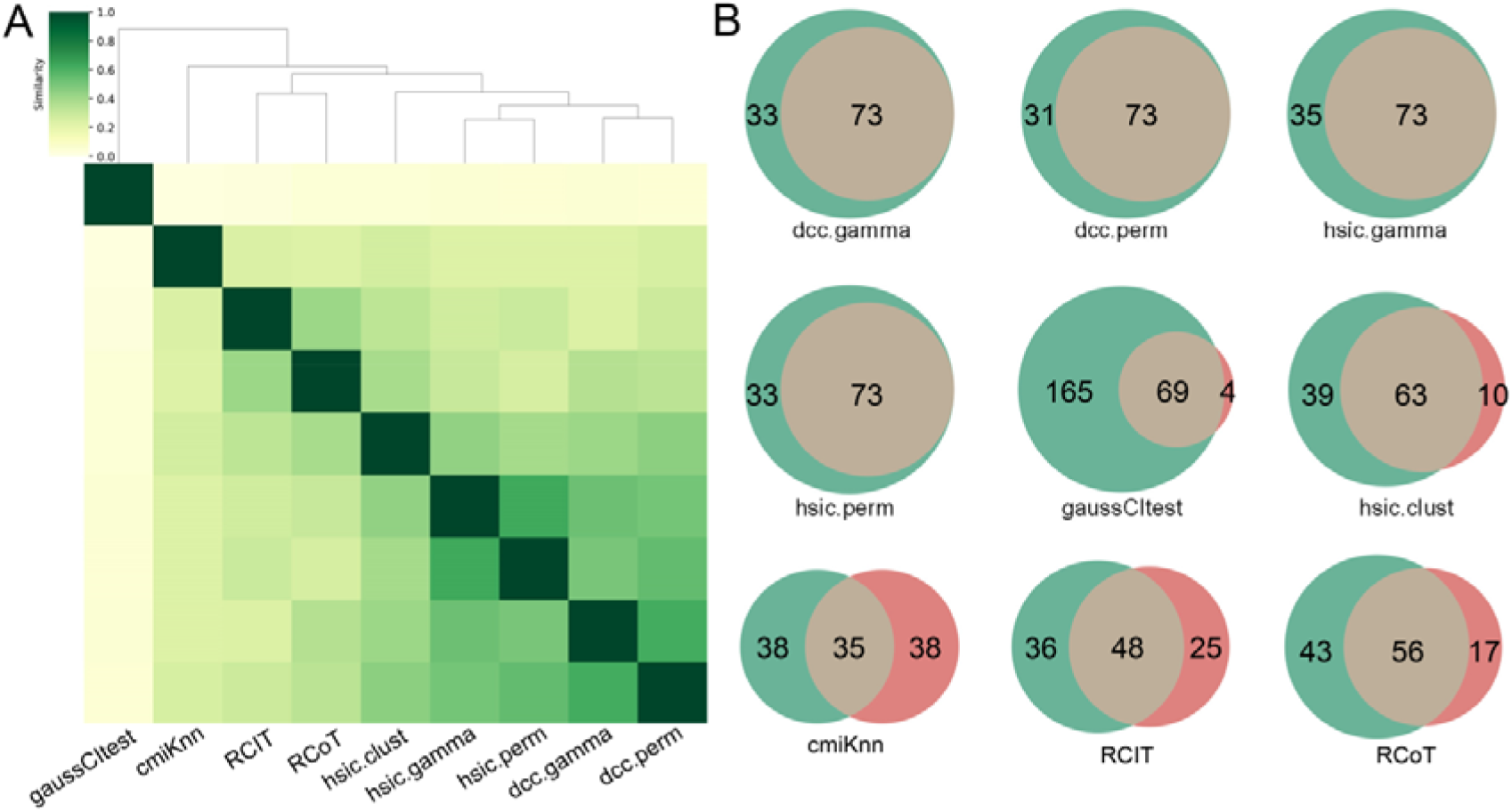
The accuracy of the 9 CI algorithms (based on 9 CI tests). (A) The cluster map measures the consistency between causal networks generated by the 9 algorithms. Darker colors indicate higher similarity, and the networks of DCC.gamma, DCC.perm, HSIC.gamma, and HSIC.perm have the highest similarity values. A consensus network built upon the four DCC and HSIC networks was used as the reference to evaluate algorithms. (B) For each algorithm’s network (green circled area), interactions overlapping the interactions in the consensus network (pink circled area) were examined. There are 73 overlapping interactions between DCC.gamma’s network and the consensus network; thus, the true positive rate of the DCC.gamma network (TPR)=73/(73+33)=68.9%. The TPR of DCC.gamma, DCC.perm, HSIC.gamma, HSIC.perm, gaussCItest, HSIC.clust, cmiKnn, RCIT and RCoT are 68.9%, 70.2%, 67.6%, 68.9%, 29.5%, 61.8%, 47.9%, 57.1%, and 56.6%.

If the scRNA-seq dataset is too large, a subset of it should be sampled. Typically, for Smart-seq2 data, 300 cells are enough, and for 10X Genomics data, 600 cells are enough. Also, HSIC.perm and DCC.perm use permutations when performing the CI test. The random sampling and permutation make causal networks inferred each time not identical. Our benchmarking and data analyses reveal that interactions inferred by DCC algorithms are highly stable (Figure 3).

**Figure 3.**
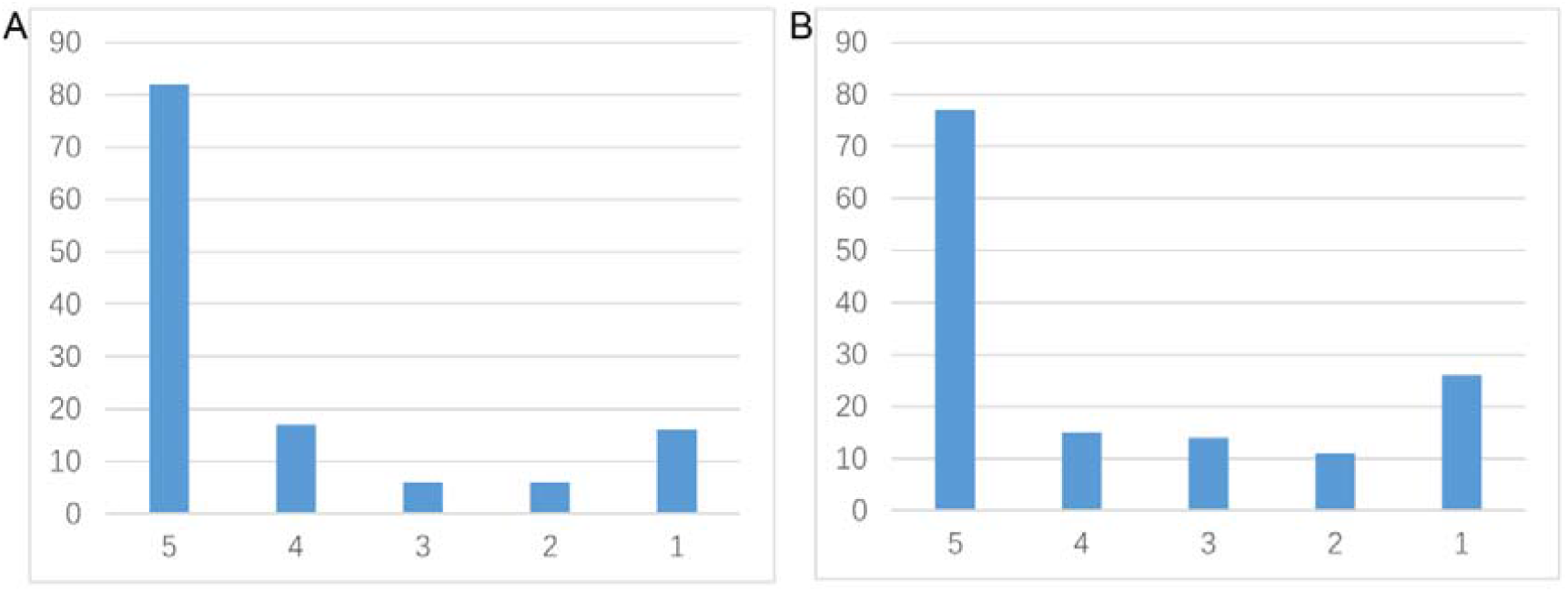
The shared and distinct interactions inferred by DCC.gamma (A) and DCC.perm (B) by running the algorithm 5 times using the dataset of lung cancer cell line H2228. 78% and 64.3% of interactions occurred stably in >=4 networks and many distinct interactions occurred in just one network, indicating that the networks inferred by the two algorithms are stable.

The following three parameters greatly influence causal discovery. “Set the alpha level” determines the statistical significance cutoff of CI test; a large alpha level causes more causal interactions to be inferred. “Select the number of cells” controls sample size; selecting more cells for causal discovery makes the inference more reliable but more time-consuming. “Select how a subset of cells is sampled” determines the way of a subset of cells is sampled. If a subset is sampled randomly, the inferred causal network is not exactly reproduable, but by running multiple times the inferred causal networks are highly consistent (Figure 3). Since each causal discovery task takes at least hours, providing an email address is necessary to make the result sent to the user automatically when it completes.

### 5. Evaluating and ensuring the reliability

A challenge for all kinds of network inferences is to verify or validate inferred networks. Inspired by using RNA spike-in to measure RNA sequencing quality, we developed a method to evaluate and ensure the reliability of causal discovery. This method includes three steps: extracting the data of some well-known genes and their interactions from some datasets as the “spike-in” data, integrating the spike-in data into the primary dataset, and applying causal discovery to the integrated dataset. In the first step, the user can pick up a spike-in data stored in the web server or design and upload a specific one; the following two steps are performed automatically. In the inferred causal network, if genes and their interactions in the spike-in data are clearly separated from genes and interactions in the primary dataset, the causal discovery should be pretty reliable (Supplementary Note 4).

### 6. Key features of different algorithms

Upon one or several response variables (i.e., genes of interest), feature selection chooses a subset of features (i.e., variables, genes) from the whole dataset by removing features unrelated or less related to response variables. A feature selection algorithm combines a search technique and an evaluation measure. After obtaining a measure between the response variable(s) and each feature, a subset of features most related to the response variable(s) is extracted. Constraint-based causal discovery algorithms identify causal relationships in a set of features in two steps: skeleton estimation (determining the skeleton of the causal network) and orientation (determining the direction of edges in the causal network). Algorithms are different in that they use different CI tests to perform the first step (the most time-consuming step). We combine the PC algorithm with 9 CI tests to form 9 causal discovery algorithms. Table 1 and Table 2 briefly describe the features and advantages/disadvantages of these feature selection and causal discovery algorithms. “+++” and “+” in the tables indicate the most and lest recommended ones.

**Table 1.**
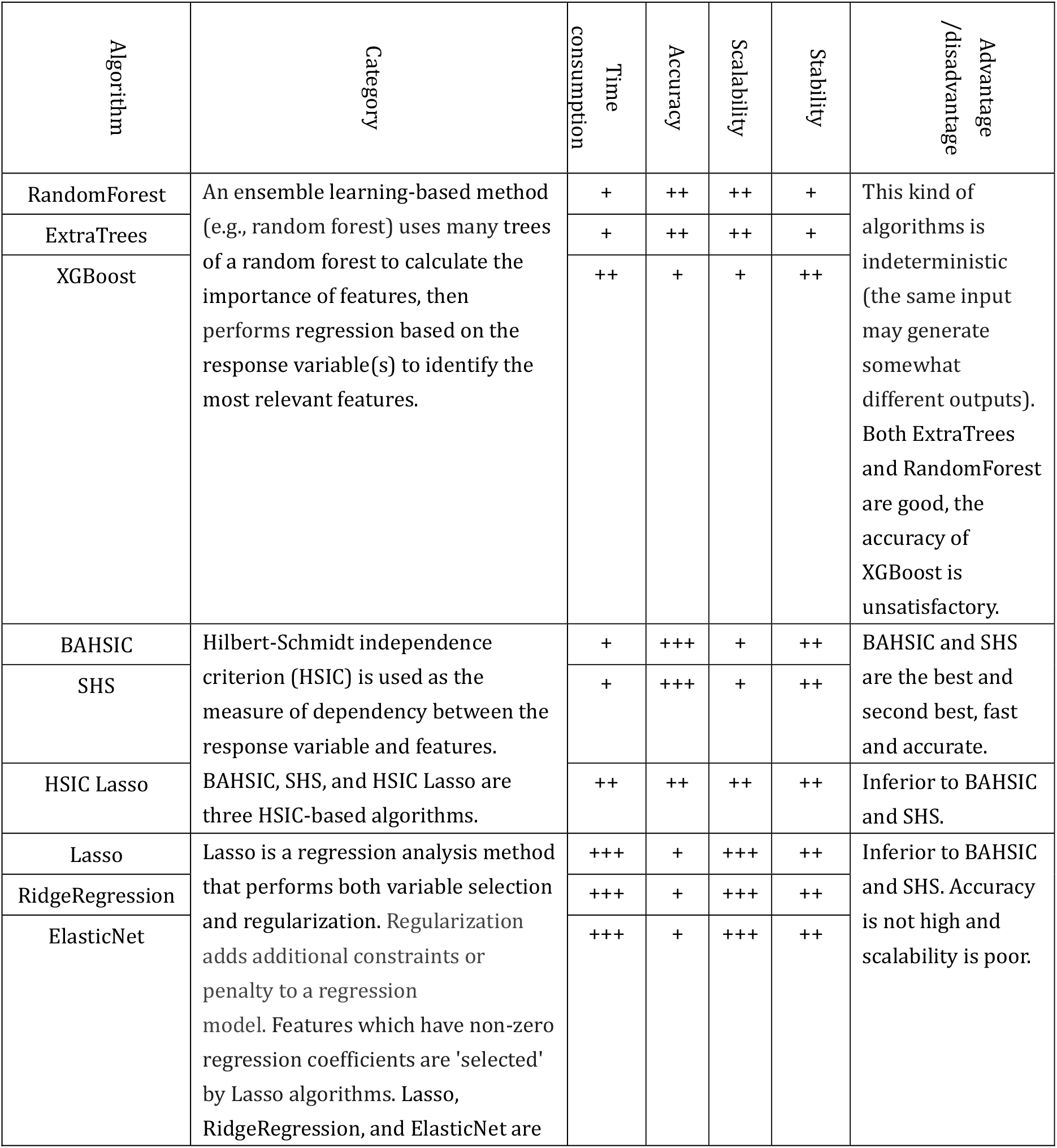
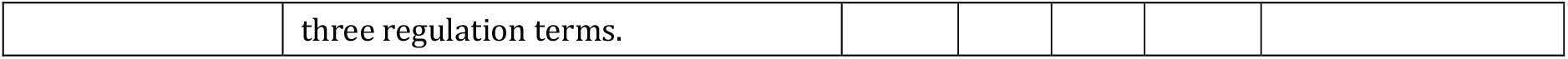
Performance of the 9 feature selection algorithms.

**Table 2.** Performance of the 9 causal discovery algorithms.

| Algorithm | Category | Time<br>consumption | Accuracy | Sample size | Stability #1 | Advantage/dis<br>advantage |
| --- | --- | --- | --- | --- | --- | --- |
| GaussCItest | Gauss CI test examines CI using partial correlation, assuming that all variables are multivariate Gaussian. This assumption impairs the performance of GussCItest, especially when data are complex. | +++ | + | +++ | +++ | Fast but inaccurate |
| CMIknn | Conditional mutual information (CMI) is a measure based on mutual information, which can be used to measure mutual dependence between two variables. | +++ | ++ | + | + | Fast but inaccurate |
| RCIT | The Kernel Conditional Independence Test (KCIT) is a powerful but time-consuming CI test. RCIT and RCoT are two approximation methods of KCIT. | ++ | ++ | ++ | ++ | Moderately accurate, fast, recommended for large-scale networks |
| RCoT |  | ++ | ++ | ++ | ++ |  |
| HSIC.clust | HSIC is a measure of dependency between two variables; HSIC(X, Y) = 0 if X and Y are unconditionally independent. HSIC.gamma and HSIC.perm employ gamma test and permutation test to estimate a *p* value. | + | ++ | ++ | ++ | Slow yet accurate, do not need large samples, recommended for small networks |
| HSIC.gamma |  | + | +++ | ++ | ++ |  |
| HSIC.perm |  | + | +++ | ++ | + |  |
| DCC.gamma | Distance covariance is an alternative to HSIC for measuring independence. DCC.gamma and DCC.perm employ Gamma test and permutation test to estimate a *p* value. | + | +++ | +++ | ++ | Slow yet most accurate, do not need large samples, recommended for small networks |
| DCC.perm |  | + | +++ | +++ | + |  |

## APPLICATIONS

### 1. The analysis of lung cancer cell lines and alveolar epithelial cells

Down-regulated MHC-II genes help cancer cells avoid being recognized by immune cells (Rooney et al., 2015); thus, identifying genes and interactions related to the down-regulation is important. To assess if causal discovery helps identify the related interactions, we examined 5 lung cancer cell lines (A549, H1975, H2228, H838, and HCC827) and the normal alveolar epithelial cells (Tian et al., 2019; Travaglini et al., 2020). For each of the six datasets, we took the 5 MHC-II genes (*HLA-DPA1, HLA-DPB1, HLA-DRA, HLA-DRB1, HLA-DRB5*) as the target genes and selected 50 feature genes (using BAHSIC, unless otherwise stated) from all genes expressed in >50% cells. Then, we applied 9 causal discovery algorithms to the 50 genes in 300 cells sampled from each of the datasets. The two DCC algorithms performed the best when processing the H2228 cells and lung alveolar epithelial cells (Figure 2; Supplementary Note 5).

Inferred networks show that down-regulated genes weakly, but up-regulated genes strongly, regulate downstream targets and that loss of activation (or inhibition) leads to down (or up) regulation. These features are biologically reasonable. Many interactions, including those among MHC-II genes and CD74, among CXCL genes, and among MHC-I genes and B2M, are supported by the STRING database (http://string-db.org) and experimental findings (Figure 4; Supplementary Fig. 12) (Castro et al., 2019; Karakikes et al., 2012; Szklarczyk et al., 2021). An interesting finding is the PRDX1→TALDO1→HSP90AA1→NQO1→PSMC4 cascade in H2228 cells. Interactions between PRDX1/TALDO1/HSP90AA1 and NQO1 were reported (Mathew et al., 2013; Yin et al., 2021), but between NQO1 and PSMC4 were not. Previous findings on NQO1 include that it determines cellular sensitivity to the antitumor agent Napabucasin in many cancer cell lines (Guo et al., 2020), is a potential poor prognostic biomarker, and is a promising therapeutic target for patients with lung cancers (Cheng et al., 2018; Siegel et al., 2012), and that mutations in *NQO1* are associated with susceptibility to various forms of cancer. Previous findings on PSMC4 include that high levels of PSMC4 (and other PSMC) transcripts were positively correlated with poor breast cancer survival (Kao et al., 2021). Thus, the inferred NQO1→PSMC4 probably somewhat explains the mechanism behind these experimental findings.

**Figure 4.**
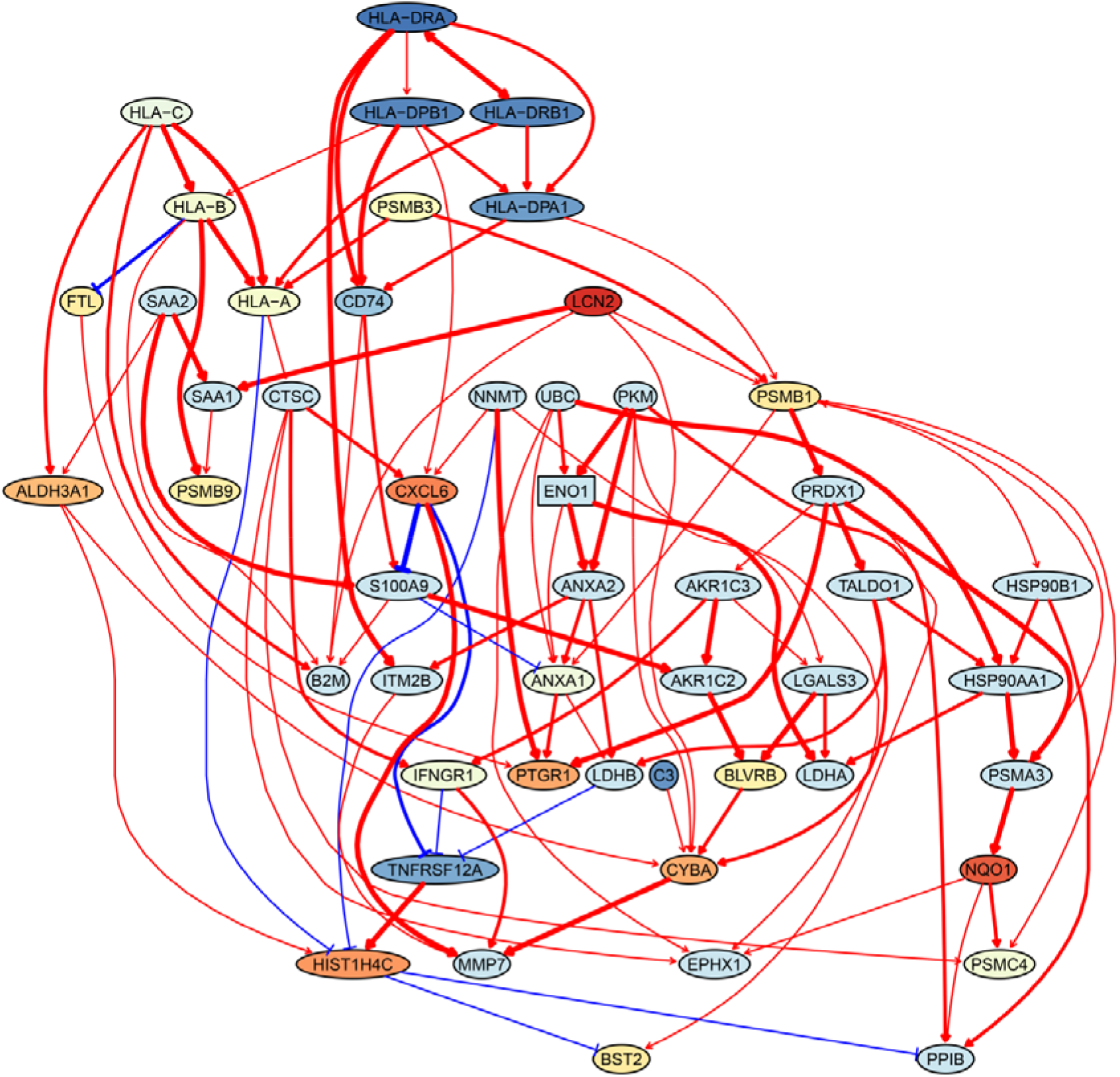
The network of the 50 genes inferred by DCC.gamma from the H2228 dataset (the alpha level for CI test was 0.1). Red → and blue -| arrows indicate activation and inhibition, and colors indicate fold changes of gene expression (from -2 to 2) compared with genes in the alveolar epithelial cells.

### 2. The analysis of macrophages isolated from glioblastoma

Macrophages critically influence glioma formation, maintenance, and progression (Gutmann, 2020), and CD74 is the master regulator of macrophage functions in glioblastoma (Alban et al., 2020; Quail and Joyce, 2017; Zeiner et al., 2015). To examine the function of CD74 in macrophages in gliomas, we used CD74 as the target gene and selected 50 genes from genes expressed in >50% macrophages isolated from glioblastoma patients (Neftel et al., 2019). In the networks of DCC algorithms (Supplementary Note 6), CD74 regulates MHC-II genes, agreeing with the finding that CD74 is an MHC-II chaperone and plays a role in the intracellular sorting of MHC class II molecules. In the network, there are interactions between C1QA/B/C, agreeing that they form the complement C1q complex. The identified TYROBP→TREM2→A2M→APOE→APOC1 cascade is supported by the reports that TREM2 is expressed in tumor macrophages in over 200 human cancer cases (Molgora et al., 2020) and that there are interactions between TREM2/A2M, TREM2/APOE, A2M/APOE, and APOE/APOC1 (Krasemann et al., 2017).

### 3. The analysis of tumor-infiltrating exhausted CD8 T cells

Tumor-infiltrating exhausted CD8 T cells are highly heterogeneous yet share common differentially expressed genes (McLane et al., 2019; Zhang et al., 2018), suggesting that CD8 T cells undergo different processes to reach exhaustion. We analyzed three exhausted CD8 T datasets isolated from human liver, colorectal, and lung cancers (Supplementary Note 7) (Guo et al., 2018; Zhang et al., 2018; Zheng et al., 2017). A key feature of CD8 T cell exhaustion identified in mice is PDCD1 upregulation by TOX (Khan et al., 2019; Scott et al., 2019; Seo et al., 2019). Using TOX and PDCD1 as the target gene, we selected 50 genes expressed in >50% exhausted CD8 T cells and 50 genes expressed in >50% non-exhausted CD8 T cells, respectively. Transcriptional regulation of PDCD1 by TOX was observed in LVMV-infected mice without mentioning any role of CXCL13 (Khan et al., 2019). Here indirect TOX→PDCD1 (via genes such as CXCL13) was inferred in exhausted CD8 cells, and direct TOX→PDCD1 was inferred in non-exhausted CD8 T cells (although the expression of TOX and PDCD1 is low in these cells) (Supplementary Figure 17). Recently, CXCL13 was found to play a critical role in T cells for effective responses to anti-PD-L1 therapies (Zhang et al., 2021). The causal discovery results help reveal differences in CD8 T cell exhaustion between species and under different pathological conditions. The PDCD1→TOX inferred in exhausted and non-exhausted CD8 T cells may indicate some feedback between TOX and PDCD1; on the proteome level, a related report is that the binding of PD1 to TOX in the cytoplasm facilitates the endocytic recycling of PD1 (Wang et al., 2019).

### 4. Identifying genes and inferring interactions that signify CD4 T cell age

How immune cells age and whether some senescence signatures reflect the aging of all cells draw wide attention (Gorgoulis et al., 2019). We analyzed gene expression in naive, TEM, rTreg, naive_Isg15, cytotoxic, and exhausted CD4 T cells from young (2-3 months, n=4) and old (22-24 months, n=4) mice (Supplementary Note 8) (Elyahu et al., 2019). For each cell type, we compared the combined data from all four young mice with the data of each old mouse to identify differentially expressed genes. If genes were expressed in >25% cells and consistently up/down-regulated (|fold change|>0) in most of the 24 comparisons, we assumed them as aging-related (Supplementary Table 4). Some of these identified genes play important roles in the aging of T cells or other cells, such as the mitochondrial genes encoding cytochrome C oxidases and the gene *Sub1* in the mTOR pathway (Bektas et al., 2019; Gorgoulis et al., 2019; Goronzy and Weyand, 2019; Walters and Cox, 2021). We directly used these genes, plus one CD4-specific biomarker (Cd28) and two reported aging biomarkers (Cdkn1b, Cdkn2d) (Gorgoulis et al., 2019; Larbi and Fulop, 2014), as feature genes to infer their interactions in different CD4 T cells in young and old mice. The causal networks unveil multiple findings (Supplementary Figure 18). First, B2m→H2-Q7 (a mouse MHC class I gene), Gm9843→Rps27rt (Gm9846), and the interactions between the five mitochondrial genes (MT-ATP6, MT-CO1/2/3, MT-Nd1) were inferred in nearly all CD4 T cells. Second, many interactions are supported by the STRING database (Supplementary Figure 13). Third, some interactions agree with experimental findings, including Sub1-|Lamtor2 (Chen et al., 2021) and the regulation of these mitochondrial genes by Lamtor2 (Morita et al., 2017). Fourth, Gm9843→Rps27rt→Junb were inferred in multiple CD4 T cells, and both Gm9843 and Rps27rt are mouse-specific. Since JUNB belongs to the AP-1 family transcription factors that are increased in all immune cells during human aging (Zheng et al., 2020), Gm9843→Rps27rt→Junb could highlight a counterpart regulation of JUNB in human immune cells.

## DISCUSSION

Various methods have been developed to infer interactions between variables from data. As surveyed recently (Nguyen et al., 2021; Pratapa et al., 2020), most methods assume linear relationships between variables and the Gaussian distribution of data. The assumptions enable these methods to run fast, capable of handling many genes or performing genome-wide predictions. Our results indicate that networks inferred by such fast methods deserve serious concern. Instead, based on kernel-based CI tests, causal discovery performs inference directly upon data observations without assuming any relationship between variables and the distribution of data (Glymour et al., 2019; Imbens and Rubin, 2015). The cost in time consumption pays off in terms of accuracy. Interacting genes and molecules within and between cells may have varied quantitative relationships, so causal discovery employing kernel-based CI tests best satisfies inferring causal interactions in varied single cells.

Several conclusions can be drawn from the benchmarking and applications. First, although kernel-based CI tests are time-consuming (Shah and Peters, 2020), applying causal discovery to a set of genes can be reasonably performed. Of note, the most time-consuming CI tests generate the most reliable results. Second, dropouts and noises in scRNA-seq data, which concern researchers and trouble correlation computation (Hou et al., 2020; Mohan and Pearl, 2018; Tu et al., 2019), can be well tolerated by kernel-based CI tests if the dataset is large enough to provide sufficient observations. Third, latent and unobserved variables influence causal discovery (just as they influence any network inference), and a solution to this problem is to evaluate whether the inference is reliable by using the “spike-in” data. Fourth, it is difficult to judge inferred interactions if without relevant information (e.g., related findings and domain knowledge).

Here are three examples showing the help of relevant information for judging inferred causal interactions. First, upon the report TOX activating PDCD1 in mice (Khan et al., 2019), whether CXCL13 is involved (or even required in humans) in the TOX-PDCD1 interaction in exhausted CD8 T cells is unclear until CXCL13 was reported to play critical roles in T cells for effective responses to anti-PD-L1 therapies (Zhang et al., 2021). Second, upon data from different cancers, inferred networks in exhausted CD8 T cells are quite different, and a recent study reports that exhausted CD8 T cells show high heterogeneity and exhaustion can follow different paths (Zheng et al., 2021). Third, it was difficult to explain the multiple genes encoding ribosomal proteins in the inferred networks in CD4 cells from old mice; a new study reports that aging impairs the ability of ribosomes to synthesize proteins efficiently (Stein et al., 2022).

### Limitations of the methods and study

First, the time consumption of the most accurate causal discovery algorithms disenables the inference of large-scale networks. Inferring multiple networks with shared genes and merging these networks into a big one is a way to circumvent this problem, but the effectiveness of the strategy remains to be confirmed. Second, it deserves noting that although time consumption pays off in accuracy, small networks could be biologically inaccurate and unreliable due to potential lack of highly related genes. Third, to make the trade-off properly between time consumption, network accuracy, and network size may need multiple rounds of trials. Fourth, the current programming language support parallel computing but does not support high-performance computing. The most time-consuming parts of the codes are to be replaced using C codes.

## TIPS FOR BEST PRACTICES

First, exploring different modules or processes needs different target genes (Figure 5). When it is unclear what gene is suitable or whether multiple genes can be co-selected, it is better to examine one by one and inspect the shared feature genes. Second, BAHSIC and SHS are the best feature selection algorithms. Third, selecting feature genes from too many candidate genes may be unreliable. Usually, filtering out some genes is necessary upon conditions such as genes are expressed in too few cells or have too low fold changes. Fourth, sometimes it is advisable to apply causal discovery to a set of genes (e.g., differentially expressed genes) without choosing a target gene and performing feature selection. Fifth, the two DCC algorithms are most recommended; it is often sufficient just to use their results to build the consensus network. Sixth, there are trade-offs between the scale, reliability, and accuracy for causal discovery. To examine many genes, using RCIT is a proper trade-off. If the dataset is large, choosing a subset of cells (e.g., 300) is a must. More cells are needed if feature genes are expressed in a small portion (e.g., 25%) of cells or if scRNA-seq data are sparse. Seventh, using a spike-in dataset and repeating causal discovery multiple rounds are two ways to ensure and improve reliability. Eighth, carefully inspect the influence of cell heterogeneity on causal discovery. Ninth, randomly sampling cells from the dataset and sampling cells with more feature genes expressed suit large and small datasets, respectively. Tenth, causal discovery identifies cell-specific causality when applied to homogeneous cells but identifies more general causality when applied to heterogeneous cells (Figure 5); in the latter case, caution is needed to interpret the results.

**Figure 5.**
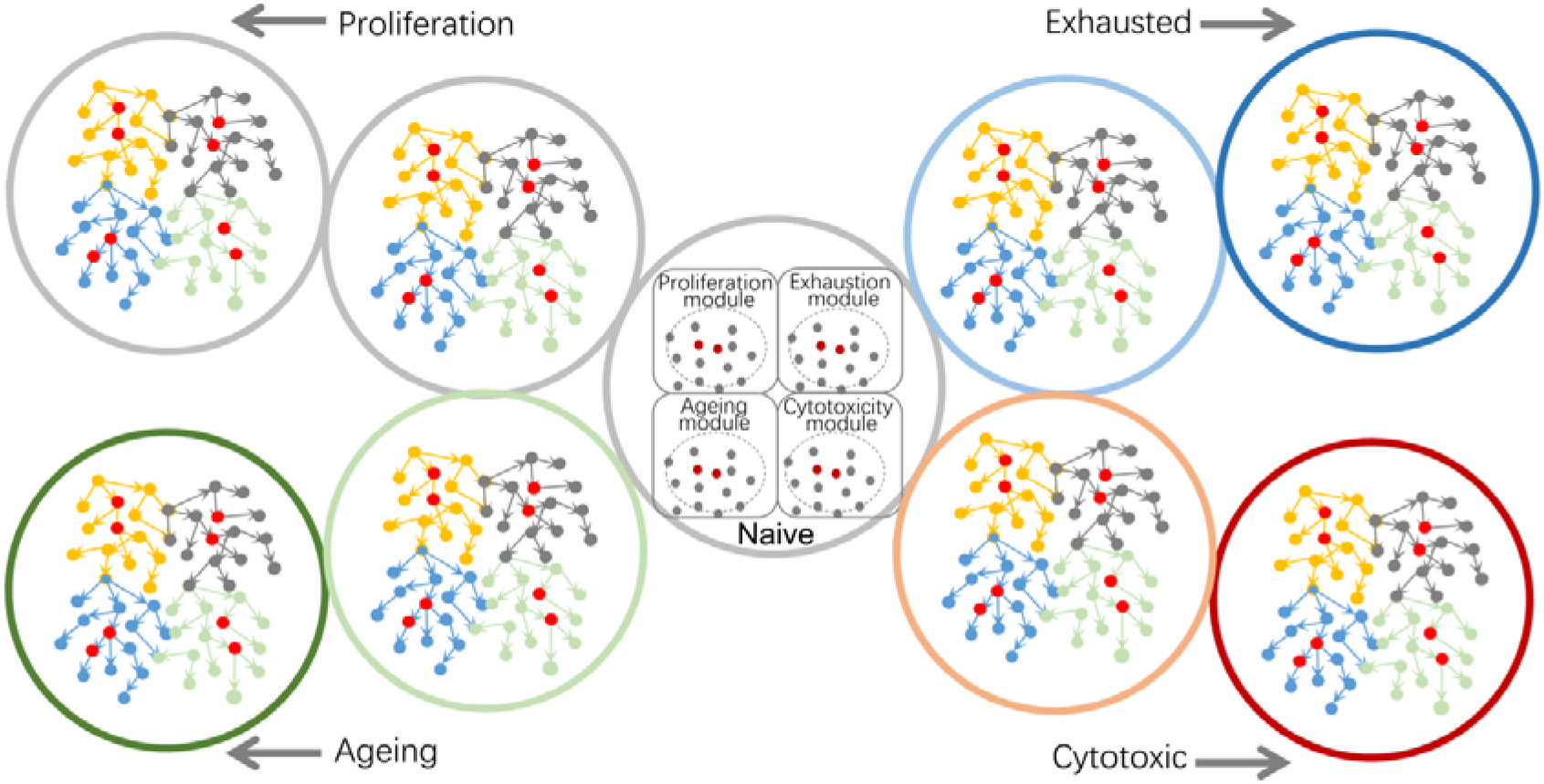
Using causal discovery to analyze different cells, cells at different stages, or different biological processes in cells. The red and grey dots within the four circles in the central cell indicate the four modules’ core genes and related genes. When exploring different biological processes, core genes in different modules should be chosen as target genes.

## Declaration of Competing Interest

The authors declare no competing interest.

## Additional information

This manuscript has one supplementary file containing supplementary tables and figures.

## Author contributions

H. Zhu designed the study and drafted the manuscript. Y.W. and J.H. performed algorithm integration and benchmarking. Y.W., H. Zhang, S.G. developed the web server. H. Zhu, A.M., Y.E., and Y.D. analyzed data. A.M. revised the manuscript. All authors have read the manuscript and consent to its publication.

## Acknowledgments

This work was supported by the National Natural Science Foundation of China (31771456) and the Department of Science and Technology of Guangdong Province (2020A1515010803). We appreciate the help from Prof Ruichu Cai at the Guangdong University of Technology.

## Data and code availability

The web server is at http://www.gaemons.net/causalcell/causalDiscovery/ (letters are capital sensitive and “http” is without ‘s’).

## Notes

### Competing Interest Statement

The authors have declared no competing interest.

